# Flagellum and toxin phase variation impacts intestinal colonization and disease development in a mouse model of Clostridioides *difficile* infection

**DOI:** 10.1101/2021.09.29.462200

**Authors:** Dominika Trzilova, Mercedes A. H. Warren, Nicole C. Gadda, Caitlin L. Williams, Rita Tamayo

**Author notes:** Corresponding Author 125 Mason Farm Rd, CB #7290, Chapel Hill, NC 27599-7290.

## Abstract

*Clostridioides difficile* is a major nosocomial pathogen that can cause severe, toxin-mediated diarrhea and pseudomembranous colitis. Recent work has shown that *C. difficile* exhibits heterogeneity in swimming motility and toxin production *in vitro* through phase variation by site-specific DNA recombination. The recombinase RecV reversibly inverts the flagellar switch sequence upstream of the *flgB* operon, leading to the ON/OFF expression of flagellum and toxin genes. How this phenomenon impacts *C. difficile* virulence *in vivo* remains unknown. We identified mutations in the right inverted repeat that reduced or prevented flagellar switch inversion by RecV. We introduced these mutations into *C. difficile* R20291 to create strains with the flagellar switch “locked” in either the ON or OFF orientation. These mutants exhibited a loss of flagellum and toxin phase variation during growth *in vitro*, yielding precisely modified mutants suitable for assessing virulence *in vivo*. In a hamster model of acute *C. difficile* infection, the phase-locked ON mutant caused greater toxin accumulation than the phase locked OFF mutant but did not differ significantly in the ability to cause acute disease symptoms. In contrast, in a mouse model, preventing flagellum and toxin phase variation affected the ability of *C. difficile* to colonize the intestinal tract and to elicit weight loss, which is attributable to differences in toxin production during infection. These results show that the ability of *C. difficile* to phase vary flagella and toxins influences colonization and disease development and suggest that the phenotypic variants generated by flagellar switch inversion have distinct capacities for causing disease.

## Introduction

Many bacterial species employ phase variation to generate phenotypic heterogeneity within a clonal population. Bacteria frequently encounter selective pressures in their environment, and phenotypic heterogeneity helps ensure survival by creating subpopulations that are differentially equipped to overcome these pressures.^1^ Phase variation typically affects the production of surface factors that directly interface with the bacterium’s environment, such as flagella, pili, and exopolysaccharides. Both mucosal pathogens and commensal species employ phase variation to balance the fitness advantages conferred by these structures with the costs of producing them; in a host environment, the ability to phase vary can promote immune evasion and persistence in the host.^2^ Phase variation can be achieved by multiple epigenetic and genetic mechanisms, including DNA modification by methylation, slipped-strand mispairing, homologous recombination, and site-specific recombination.^1, 3^

In many pathogens including *Acinetobacter baumannii, Bordetella bronchiseptica*, and *Streptococcus pneumoniae*, host selective pressures can substantially affect the composition of a phase-variable bacterial population during infection.^4-11^ All three of these pathogens can form distinct subpopulations differentially equipped for survival in disparate environments. However, determining the importance of phase variation (rather than the phase-variable trait) in pathogenesis can be challenging. Studies in which phase variation has been eliminated to create “phase-locked” mutants have been valuable for determining the impact of phase variation itself. In *B. bronchiseptica*, phase-locked Bvg+ and Bvg-strains were created by either deleting *bvgS* encoding the sensor kinase in the BvgAS system (Bvg-phase) or mutating BvgS to be constitutively active (Bvg+ phase).^4^ Phase-locked strains have also been created by mutating site-specific recombinases that mediate inversion of one or multiple loci, e.g., Mpi modulating polysaccharide production in *Bacteroides fragilis*.^12^ Finally, in uropathogenic *Escherichia coli* (UPEC), mutation of inverted repeats critical for site-specific recombination generated populations unable to phase vary the production of fimbriae.^13, 14^ These studies led to improved understanding of host-microbe interactions and the importance of phase variation in fitness of bacterial pathogens in diverse host environments.

*Clostridioides difficile* is a gram-positive, spore-forming anaerobe that is currently the leading cause of antibiotic-associated diarrheal disease and one of the most common causes of nosocomial infection. *C. difficile* infection (CDI) is primarily mediated by two toxins, TcdA and TcdB, that glucosylate and inactivate Rho family GTPases leading to perturbation of the actin cytoskeleton.^15-17^ During infection, TcdA and TcdB disrupt the intestinal epithelial barrier resulting in inflammation, immune cell recruitment, and development of diarrheal symptoms,^18^ and evidence suggests both toxins are important for disease development in animal models.^19-21^

Several recent studies have shown that *C. difficile* exhibits substantial phenotypic heterogeneity via phase variation by site-specific DNA recombination.^22-26^ This mechanism of phase variation is mediated by serine or tyrosine DNA recombinases that recognize sequences containing inverted repeats and catalyze strand exchange, leading to inversion of the intervening DNA sequence.^27, 28^ Eight DNA sequences that can undergo inversion have been identified in *C. difficile*, though individual strains may contain only a subset.^24, 29^ Three of these sequences have been experimentally demonstrated to modulate expression of adjacent genes leading to phase variation of the encoded factors: the cell wall protein CwpV,^22, 30^ flagella,^23, 31^ and the CmrRST signal transduction system.^26^

*C. difficile* flagella are required for swimming motility and contribute to adherence to intestinal epithelial cells, colonization, and virulence in animal models of infection.^32-35^ Flagellar genes in *C. difficile* are organized in multiple operons that are expressed in a hierarchical manner coordinated by the sigma factor SigD (also known as FliA and σ ^28^).^36-39^ SigD also promotes expression of toxin genes by activating transcription of *tcdR*, which encodes a direct positive regulator of *tcdA* and *tcdB*, linking toxin production to flagellar gene transcription.^38-40^ Consistent with these findings, phase variation of flagella results in concomitant phase variation of toxin biosynthesis *in vitro*.^23, 25, 41^ This phase variation occurs as a result of inversion of a DNA sequence, termed the flagellar (*flg*) switch, upstream of the *flgB* coding sequence and mapping to the 5’ untranslated region of the *flgB* operon.^23^ The flagellar switch is flanked by imperfect inverted repeats and contains regulatory features that control gene expression by a mechanism dependent on Rho-mediated transcription termination.^41^ This process generates a phenotypically heterogeneous population consisting of *flg* ON cells that are motile and toxigenic and *flg* OFF cells that are aflagellate, nonmotile, and attenuated for toxin production.^23, 41^

The roles of flagella and toxins in *C. difficile* pathogenesis are well studied, but the importance of the heterogeneity generated by their phase variation to disease development is unknown. The goal of this study was to analyze the effects of preventing phase variation of the flagellar switch on *C. difficile* physiology and virulence. Phase-locked ‘ON’ and ‘OFF’ mutants were previously generated by eliminating the site-specific tyrosine recombinase, RecV, and identifying isolates with the flagellar switch locked in either orientation.^22^ However, RecV is required for inversion of three other sequences (upstream of *cwpV, cmrRST*, and CDR20291_0963) and influences inversion of two additional sequences (upstream of CDR20291_0685 and CDR20291_1514).^22-24, 26, 42^ The broad regulation exerted by RecV thus results in pleiotropic effects that limit the utility of *recV* mutants for studying the impact of flagellum and toxin phase variation. To circumvent these drawbacks, in this study we instead created phase-locked strains by mutating residues in the right inverted repeat predicted to be critical for flagellar switch inversion. We identified mutations that cause a partial or full attenuation of flagellar switch inversion, and *in vitro* characterization of these mutants showed a corresponding reduction or loss of flagellum and toxin phase variation. The locked ON and OFF mutants were assessed in hamster and mouse models of CDI for altered colonization, virulence properties, and toxin production. In a hamster model of acute CDI, the phase-locked mutants did not differ from wildtype in ability to cause disease and showed a modest defect in colonization, despite significant differences in toxin accumulation in the cecum between the locked-ON and - OFF mutants. In contrast, in a mouse model of CDI, the locked-ON mutant elicited significantly greater weight loss and maintained higher colonization levels compared to wildtype and the locked-OFF mutant. Differences were attributable at least in part to toxin levels achieved by these strains during infection. These results indicate that the capacity of *C. difficile* to phase vary flagellum and toxin biosynthesis during infection impacts its ability to colonize and cause disease *in vivo*.

## Results

### Identification of nucleotides in the right inverted repeat of the flagellar switch that are important for inversion

To determine the role of flagellum and toxin phase variation in *C. difficile* infection, we aimed to create mutants incapable of inverting the flagellar switch while avoiding the pleiotropic effects of inactivating *recV*. We focused on the flagellar switch inverted repeats (IRs), as these regions are typically directly recognized by the site-specific recombinase and important for switch inversion. To identify candidate nucleotides in the IRs required for inversion of the flagellar switch by RecV, we performed an alignment of the IRs of the six RecV-invertible sequences in *C. difficile*. Although the invertible sequences differ in length and nucleotide sequence, the IRs share some sequence similarity suggesting that specific residues are important for interacting with RecV. Sequence conservation is strongest among the left inverted repeats (LIRs) of five invertible sequences and right inverted repeat (RIR) of the *cwpV* switch; lower identity is apparent among the respective RIRs (and LIR of *cwpV*) (Figure 1A).

**Figure 1.**
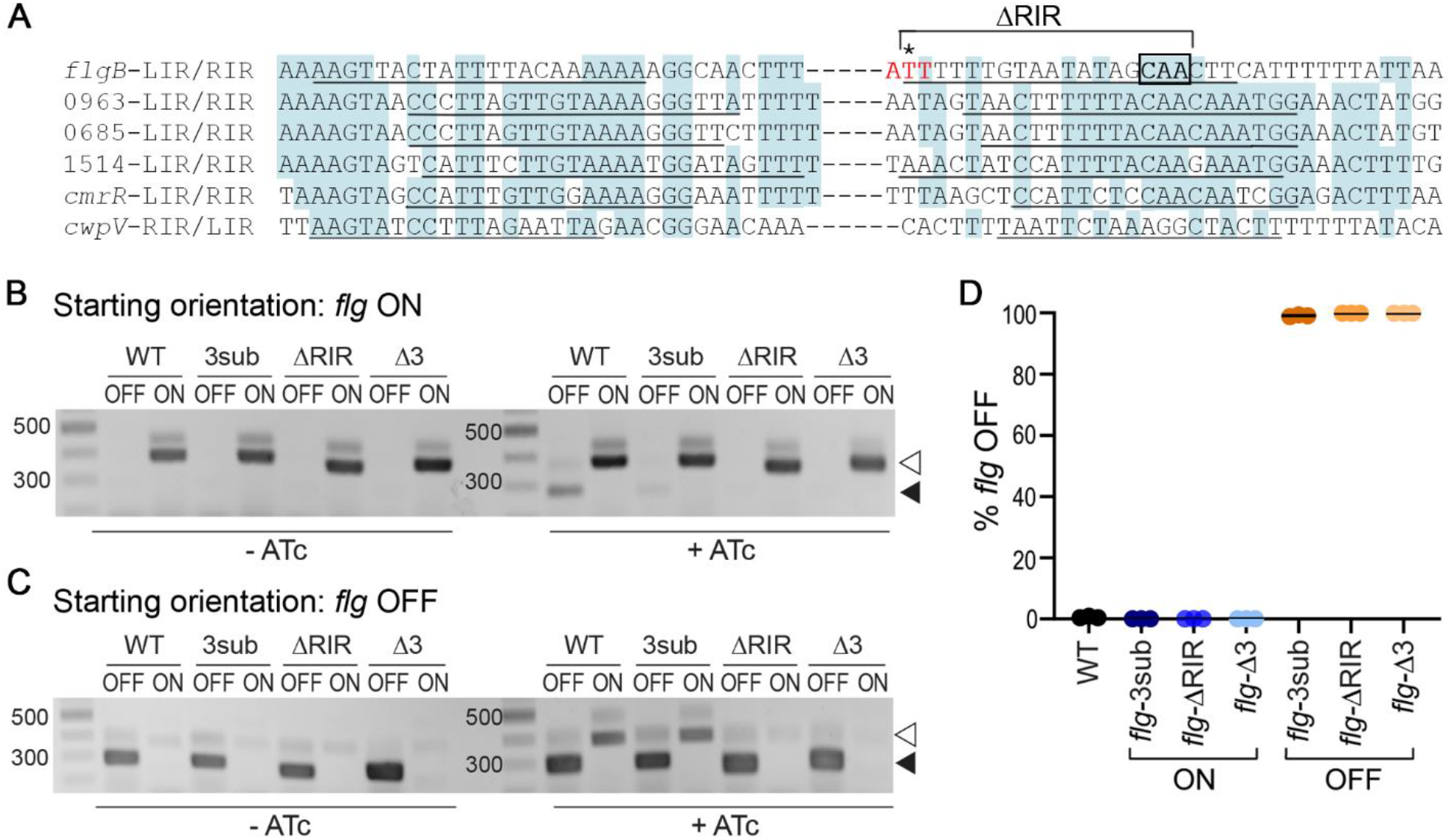
Mutations in *flg* RIR affect inversion in *E. coli* and *C. difficile*. A. Alignment of inverted repeats flanking the invertible DNA sequences affected by RecV. Shading denotes residues conserved in at least 4 of the 6 repeats. Putative inverted repeats are underlined. Hyphens between LIRs and RIRs represent the intervening sequences, which vary in length and nucleotide sequence. For the *flgB*-RIR, the site of cleavage by RecV is indicated with an asterisk. This nucleotide and the two adjacent residues, indicated in red, were deleted in *flg-*Δ3 constructs/strains. Note that the adenine 5’ of the cleavage site is present in the *flg* ON sequence, whereas a thymine is present in *flg* OFF; constructs for mutagenesis were created in both *flg* orientations. The conserved CAA nucleotides mutated in *flg*-3sub constructs/strains are boxed in black. Nucleotides deleted in *flg*-ΔRIR bacteria/constructs are indicated. LIR, RIR = left, right inverted repeats. Numbers indicate locus tags in *C. difficile* R20291. B, C. Orientation-specific PCR to examine flagellar switch inversion in *E. coli* bearing wild-type or mutated inverted repeat target sequences. The starting orientation of the flagellar switch is indicated: *flg* ON (B) or *flg* OFF (C). Absence or presence of ATc for induction of *recV* expression is shown (-ATc/+ATc). The *flg* ON and *flg* OFF products are indicated with white and black arrows, respectively. Shown are representative images of three independent experiments. (D) Analysis of flagellar switch orientation in *C. difficile* RIR mutants by quantitative orientation-specific PCR. Means and standard deviations for three biological replicates are shown.

We recently showed that the flagellar switch impacts gene expression in a mechanism acting after transcription initiation and occurring within the leader region of the *flgB* operon mRNA. While not fully elucidated, this regulation is dependent on Rho-mediated transcription termination that preferentially impacts *flg* OFF mRNA, a mechanism that requires Rho to interact with the mRNA either within or upstream of the flagellar switch.^41^ To minimize the risk of interfering with Rho-mediated regulation, we chose to mutate the RIR downstream of the flagellar switch.

Because the starting orientation of an invertible element can impact the efficiency of inversion by a recombinase, both *flg* ON and *flg* OFF versions were created for each mutation. Three sites were selected for mutagenesis. First, we deleted 18 of the 21 bp of the *flg* RIR to create *flg-*ΔRIR ON and OFF mutant sequences, which we anticipated would prevent switch inversion;^43^ however, the larger deletion presents greater risk of altering Rho-mediated regulation. Second, we chose three highly conserved CAA nucleotides in the *flg* RIR for substitution with GTT to create *flg-*3sub ON and OFF mutant sequences (Figure 1A). Third, we targeted the previously identified nucleotide where the DNA strand is cleaved by RecV to catalyze strand exchange (Figure 1A).^24^ This residue and the two adjacent nucleotides were deleted to create *flg-*Δ3 ON and OFF mutant sequences.

Due to the challenges of creating unmarked chromosomal mutations in *C. difficile*, we first evaluated the effects of the RIR mutations on flagellar switch inversion using a previously described method employing *E. coli* as a heterologous host.^22, 23^ In this assay, the *E. coli* strains bear two plasmids. One plasmid contains *recV* under the control of an anhydrotetracycline (ATc)-inducible promoter, and the other plasmid contains the target flagellar switch sequence.^23^ Primers specific to each orientation of the switch are then used for detection by PCR. Each of these plasmid-borne target sequences was co-transformed into *E. coli* with pRecV, and the resulting strains were grown with or without ATc to induce *recV* expression then subjected to PCR with orientation-specific primers. As observed previously, only the starting orientation of the flagellar switch sequence was detected in the absence of *recV* induction with ATc for both *flg* ON and *flg* OFF constructs (Figure 1B, C, left). Upon induction with ATc, inversion of the wild-type flagellar switch sequence was detected, evident by the appearance of the *flg* OFF product from the *flg* ON target (Figure 1B, right) and the *flg* ON product from the *flg* OFF target (Figure 1C, right). The *flg*-3sub mutation appeared to reduce but not eliminate inversion by RecV, as the product for the inverted sequence remained detectable (Figure 1B, C, right). In contrast, no inversion was detected for the *flg*-ΔRIR and *flg*-Δ3 target sequences, regardless of the starting orientation of the flagellar switch (Figure 1B, C). The starting orientation of the flagellar switch did not affect invertibility of any of the target sequences by RecV in this assay. These data suggest that the *flg*-ΔRIR and *flg*-Δ3 mutations in the RIR render the flagellar switch into a “locked” state, while the *flg*-3sub mutation impairs flagellar switch inversion.

### Loss of flagellar switch inversion in the RIR mutants leads to *C. difficile* phase-locked for motility and toxin production

We next sought to determine the effects of the RIR mutations on flagellar switch inversion and phase variation in *C. difficile*. We used allelic exchange to create six R20291 mutant strains: *flg-*3sub ON, *flg*-ΔRIR ON, *flg*-Δ3 ON, *flg-*3sub OFF, *flg*-ΔRIR OFF, and *flg*-Δ3 OFF. The process of generating strains with these precise mutations was facilitated by first deleting the 5’UTR region, then restoring that region with the desired nucleotide changes incorporated. These mutants were confirmed to have the expected flagellar switch orientation using quantitative PCR with orientation-specific primers (OS-qPCR) (Figure 1D). Each mutant contained the flagellar switch exclusively in the anticipated ON or OFF orientation (0 ± 0 % *flg* OFF for the *flg* ON mutants, 100 ± 0 % *flg* OFF for the *flg* OFF mutants) after growth in rich, liquid medium, in contrast to wildtype which exhibited heterogeneity (0.6 ± 0.3 % *flg* OFF).

To establish that these genetically locked mutants are also phenotypically locked, we first tested these strains in soft agar swimming motility assays. We found that, as expected, the wildtype and *flg* ON mutant strains (*flg*-ΔRIR, *flg*-Δ3, and *flg*-3sub) exhibited comparable motility (Figure 2A, B). In contrast, examination of the equivalent mutations in the *flg* OFF background revealed distinct effects of the mutations on flagellar phase variation. The *flg*-ΔRIR OFF and *flg-*Δ3 OFF mutants remained non-motile, equal to the nonmotile *sigD-*null control, indicating that these mutations prevent phase variation. However, the *flg*-3sub OFF mutant exhibited motility in this assay. These results are consistent with the data in Figure 1 indicating that the 3sub mutation reduces but does not eliminate flagellar switch inversion, and they suggest that the motility medium presented a selective pressure for the *flg* ON variants.

**Figure 2.**
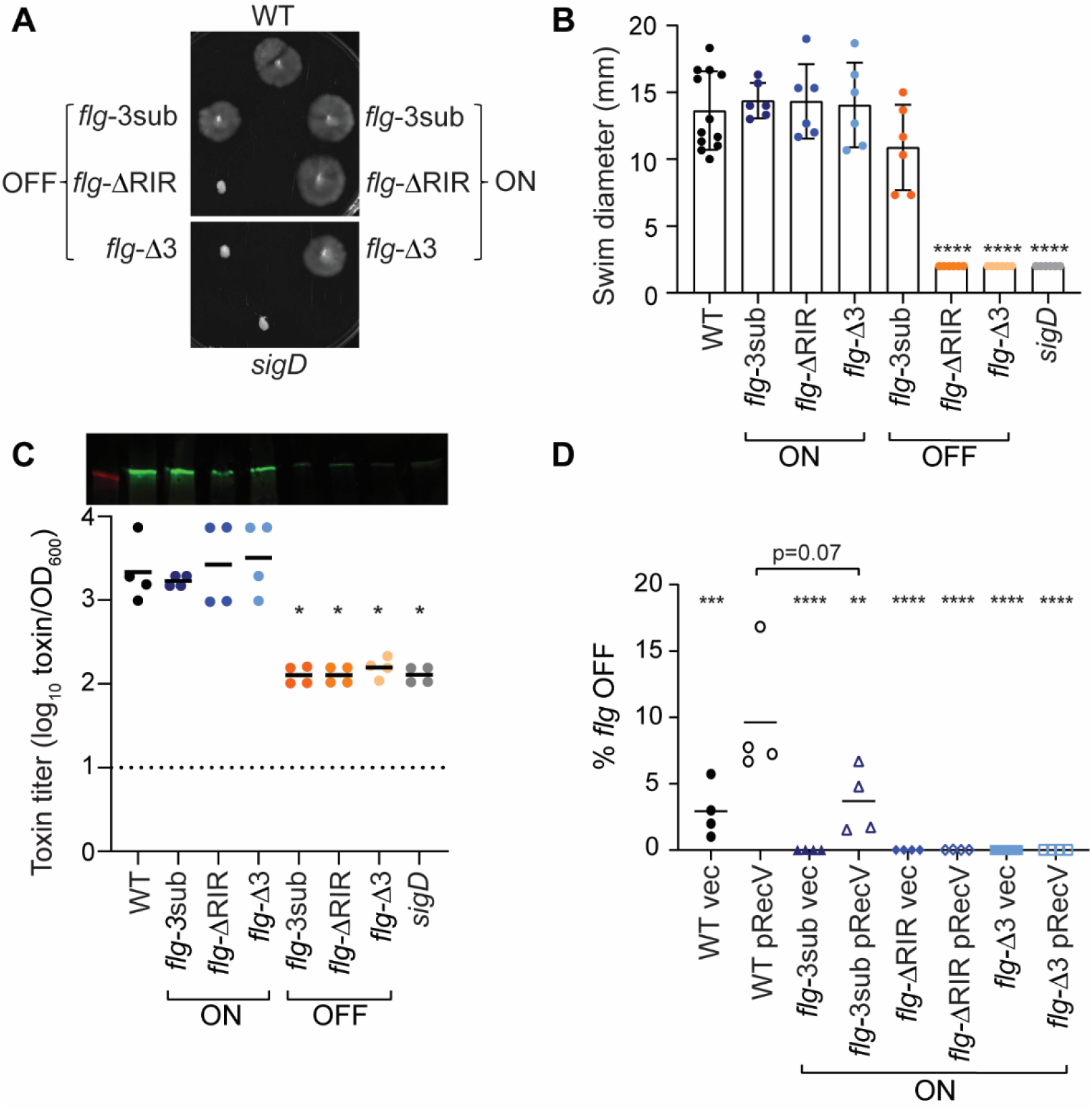
Mutations in *flg* RIR affect *C. difficile* motility and toxin production. (A) Representative image of swimming motility in soft agar medium of *C. difficile* R20291 (WT), *flg-*3sub ON and OFF, *flg-*ΔRIR ON and OFF, and *flg-*Δ3 ON and OFF, and *sigD*-null non-motile control, incubated for 48 hours. (B) Quantification of swimming motility after 48 h of strains in (A). (C) Immunoblot detection of TcdA and toxin titers after growth in TY broth. For immunoblot, a representative image of three independent experiments is shown. Toxin titers of supernatants from overnight bacterial cultures were calculated as the reciprocal of the highest dilution that causes ≥80% rounding of Vero cells, expressed after log-transformation and normalization to OD_600_ of the cultures. (B, C) Each symbol represents one biological replicate, and dotted line represents the limit of detection. \**p*<0.05 by one-way ANOVA with Dunnett’s post-test comparing values to *flg-*Δ3 ON. (D) Quantitative orientation-specific PCR of the flagellar switch in WT, *flg-*3sub ON, *flg-*ΔRIR ON, and *flg-*Δ3 ON mutants expressing *recV* (pRecV) or bearing vector. Means and standard deviations are shown. \*\*\*\**p<*0.0001, \*\*\**p*<0.001, \*\**p*<0.01 by one-way ANOVA and Dunnett’s post-test comparing values to WT pRecV. P value for comparison of WT pRecV and *flg*-3sub ON pRecV was determined by unpaired two-tailed t-test. (B-D) Means and standard deviations are shown.

Because toxin gene expression is linked to transcription of the *flgB* operon via SigD,^38, 39^ we evaluated toxin production in the *flg* RIR mutant strains. By immunoblot, the *flg*-ΔRIR, *flg-*Δ3, and *flg-*3sub ON strains produced TcdA at levels equivalent to the wildtype parent after growth in TY broth (Figure 2C). TcdA was undetectable in the three *flg* OFF mutants, similar to the *sigD*-null control. Using a Vero cell rounding assay, which detects the activities of TcdA and TcdB, to quantify the toxin produced by these strains in broth culture, we found that the three *flg* OFF mutants and *sigD* control cultures contained significantly lower toxin titers than the *flg* ON mutants and wildtype (Figure 2C). Therefore, mutations in *flg* RIR that impede inversion of the flagellar switch concomitantly impact toxin production *in vitro*.

While a swimming motility assay can show the lack of flagellar switch inversion from OFF to ON in non-motile bacteria, it cannot detect inversion from ON to OFF in motile bacteria (including the *flg* ON RIR mutants) because the motile phenotype dominates in this assay. As an alternative way to determine whether the *flg*-3sub, *flg*-ΔRIR and *flg*-Δ3 ON mutants are capable of inversion, we assessed the effect of *recV* overexpression on flagellar switch inversion from ON to OFF using qPCR with orientation-specific primers (OS-qPCR). As seen previously, the wild-type R20291 populations were skewed toward the *flg* ON orientation, with less than 5% *flg* OFF cells, and expression of *recV* increased *flg* switch inversion resulting in a larger *flg* OFF population (Figure 2D).^23^ The three *flg* ON RIR mutants bearing vector, with *recV* expressed at its natural levels, consisted of only *flg* ON bacteria (0% *flg* OFF, n=4). Despite *recV* overexpression, no *flg* OFF bacteria were detected in the *flg*-ΔRIR and *flg*-Δ3 ON mutants. However, overexpression of *recV* increased the subpopulation with the OFF orientation in the *flg*-3sub ON mutant, though potentially at a lower frequency than in the wildtype. Together with the *flg* OFF motility data and the *E. coli* inversion experiments, these results show that deletion of the RIR (ΔRIR) or three nucleotides at the site of recombination (Δ3) results in complete loss of flagellar switch inversion and phase variation, while a substitution of three conserved residues (3sub) reduces the frequency of inversion.

### Preventing flagellar switch inversion affects colonization and toxin accumulation in a hamster model of infection

*In vitro*, restricting inversion of the flagellar switch affects both motility and toxin production. Because both characteristics are important during CDI, we analyzed the virulence of *flg* RIR mutants in a hamster model of infection, which is particularly sensitive to TcdA and TcdB and manifests acute CDI.^19, 20, 44^ Prior work showed that inactivation of genes in the *flgB* operon, which resulted in reduced toxin production, also attenuated virulence in this model,^32, 33^ so we anticipated that the hamster model would distinguish the virulence of phase-locked *flg* ON and *flg* OFF mutants. To conserve animals, we narrowed this study to the *flg*-Δ3 mutants which contain the smallest mutation that prevents flagellum and toxin phase variation. Antibiotic-treated male and female Syrian golden hamsters were inoculated with 1000 spores of wild-type R20291, *flg-*Δ3 ON, or *flg-*Δ3 OFF. These strains had no differences in growth (Figure S1), germination (Figure S2), or sporulation (Figure S3) *in vitro*. The animals were monitored for disease symptoms, including diarrhea and weight loss, and were euthanized if they exhibited hallmarks of disease as detailed in Materials and Methods. Initially, hamsters infected with the *flg-*Δ3 ON or *flg-*Δ3 OFF mutants appeared to become acutely symptomatic sooner than those infected with wildtype, however, there were no significant differences in time to euthanasia for animals infected with any of the strains (Figure 3A).

**Figure 3.**
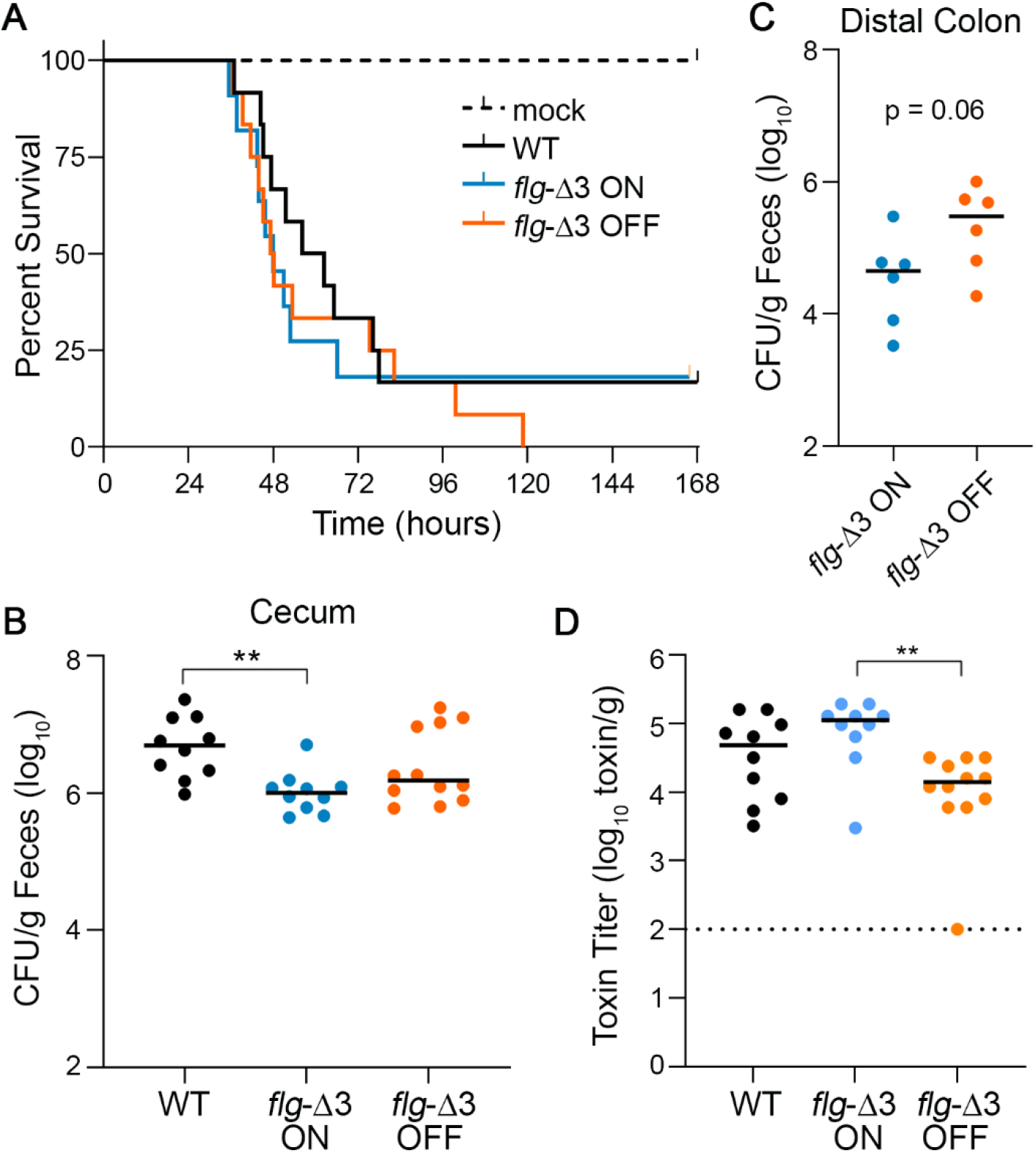
Interfering with flagellar switch inversion affects toxin accumulation and bacterial burden in a hamster model of CDI. Antibiotic-treated male and female Syrian Golden hamsters were inoculated with 1000 spores of wild-type R20291 (WT), *flg*-Δ3 ON, and *flg-*Δ3 OFF. Mock-inoculated animals were included in each experiment. Data are combined from two independent experiments testing strains in 3 male and 3 female hamsters, for 12 total hamsters per strain. (A) Kaplan-Meier analysis of survival. (B) CFU in cecal contents. \*\**p*<0.01 by Kruskal-Wallis test with Dunn’s post-test. (C) CFU from homogenized distal colon from six animals of one experiment. P value was determined by Mann-Whitney test. (D) Toxin titers in cecal contents calculated as the reciprocal of the highest dilution to cause ≥80% rounding of Vero cells. No cell rounding occurred when treated with diluted cecal contents from mock-inoculated animals. Bars indicate the means; dotted line represents the limit of detection. \*\**p<*0.01 with Kruskal-Wallis test with Dunn’s post-test. (B, C, D) Symbols indicate CFU from individual animals and bars indicate medians.

To determine bacterial burden, cecal contents collected immediately after euthanasia were serially diluted and plated on taurocholate cycloserine cefoxitin fructose agar (TCCFA) to enumerate *C. difficile* colony forming units (CFU). All animals that succumbed to disease had *C. difficile* detectable in their cecal contents (10^4^ CFU/g to 10^7^ CFU/g). Wild-type R20291 was present in 5.3-fold greater CFU compared to the *flg*-Δ3 ON mutant (p < 0.01); there were no significant differences among the other strains (Figure 3B). In the distal colons there was a similar trend, with 5-fold more CFU/g for the *flg*-Δ3 OFF mutant compared to the *flg*-Δ3 ON mutant (p *=* 0.06) (Figure 3C). These results suggest that flagellar phase-locked ON mutants are less fit than the phase-locked OFF mutants in the hamster intestinal tract, though the effect was modest.

Because of the link between production of flagella and toxins *in vitro*, we analyzed toxin titers in cecal contents of hamsters that succumbed to disease using the Vero cell rounding assay.^45^ All samples from infected animals caused detectable cell rounding, while no cell rounding occurred when treated with diluted cecal contents from mock-inoculated animals. The toxin titers for the *flg*-Δ3 ON samples were 6.5-fold higher compared to the *flg*-Δ3 OFF (p < 0.01) (Figure 3D). These results indicate that, despite the lack of difference in ability to cause acute CDI, preventing phase variation led to significant differences in toxin accumulation during infection, and flagellum and toxin gene expression are linked during infection as observed *in vitro*.

### Preventing flagellar switch inversion impacts colonization and disease dynamics of *C. difficile* in a mouse model

While the hamster models acute CDI, the mouse typically develops less severe disease and serves as a model of *C. difficile* colonization. Antibiotic-treated male and female C57BL/6 mice were inoculated with 10,000 spores of wild-type R20291, *flg-*Δ3 ON, or *flg-*Δ3 OFF.^46^ Over the following 10 days, the mice were monitored for diarrhea and weight loss, and fecal samples were collected daily to assess bacterial burden and toxin levels. Bacterial burden in feces achieved the highest levels between days 1 and 4 post-inoculation (p.i.) (Figures 4A, S4). Although all three strains were present in equivalent numbers during this time frame, the *flg-*Δ3 ON elicited significantly greater weight loss than *flg-*Δ3 OFF on days 1 through 3 (Figures 4C, S5). Wildtype resulted in intermediate weight loss, with significant differences from *flg-*Δ3 ON and *flg-*Δ3 OFF on days 2 and 3 p.i. (Figure 4D). The WT- and *flg-*Δ3 ON-infected mice began recovering weight on day 3 p.i., with no significant differences in weight loss among the groups on day 4 and later. On day 4 p.i., the CFU/g feces from wildtype and *flg*-Δ3 OFF infected mice began to decline, often to below the limit of detection, while *flg-*Δ3 ON showed a comparatively modest decline between days 3 and 4 and then was maintained at ∼10^5^ CFU/g feces in all mice through the duration of the experiment (Figures 4A, 4B, S4).^41^ The *flg-*Δ3 ON mutant was present in significantly higher numbers than wildtype on days 6 through 10 p.i. (Figures 4A, S4). The *flg-*Δ3 OFF mutant colonized to levels similar to wildtype or intermediate between wildtype and *flg-*Δ3 ON.

**Figure 4.**
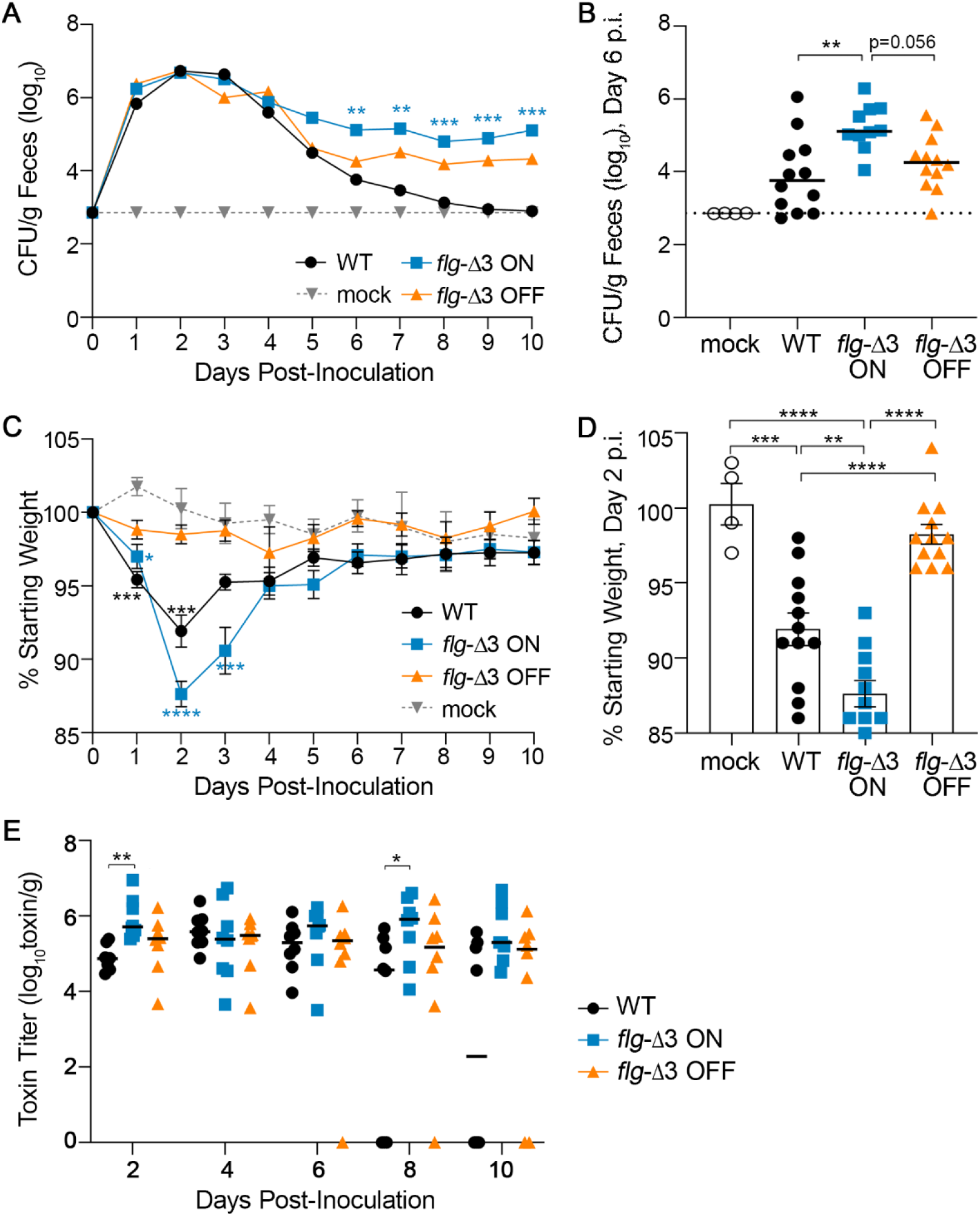
Locking the flagellar switch in the ON orientation exacerbates disease and increases persistence of *C. difficile* in a mouse model of CDI. Antibiotic-treated male and female C57BL/6 mice were inoculated with 100,000 spores of wild-type R20291 (WT), *flg*-Δ3 ON, and *flg-*Δ3 OFF. Mock-inoculated animals were included in each experiment. Data are combined from two independent experiments testing strains in 3 male and 3 female mice, for 12 total mice per strain. (A) CFU enumerated in fecal samples collected every 24 hours post-inoculation (p.i.). Asterisks indicate statistical comparison to WT data at that time point. (B) CFU per gram feces collected on day 6 p.i.; data for days 5 - 10 shown in Figure S4. Symbols in each group distinguish results from two independent experiments. (C) Animal weights determined every 24 hours post-inoculation, expressed as a percentage of the mouse’s weight at day 0. Asterisks indicate statistical comparison to mock data at that time point. (D) Animal weights at Day 2 p.i.; data for days 1 - 6 shown in Figure S5. (E) Toxin titers in fecal samples calculated as the reciprocal of the highest dilution to cause ≥80% rounding of Vero cells. No cell rounding occurred when treated with diluted fecal contents from mock-inoculated animals. (A-E) Symbols indicate values from individual animals; dotted lines represent limit of detection. (A, B, E) Bars indicate the medians. Statistical significance was determined using the Kruskal-Wallis test and Dunn’s post-test. (C, D) Bars indicate means and standard error. Statistical significance was determined by one-way ANOVA with Tukey’s post-test. \**p*<0.05, \*\**p<*0.01, \*\*\**p*<0.001, \*\*\*\**p*<0.0001.

We additionally ensured that *flg-*Δ3 ON and *flg-*Δ3 OFF remained phase-locked by evaluating them at the infection endpoint. OS-qPCR analysis of genomic DNA purified from fecal samples indicated that the flagellar switch remained locked in *flg-*Δ3 ON (collected on day 9 p.i.; data not shown). The lower colonization levels by *flg-*Δ3 OFF after day 5 precluded OS-qPCR analysis. As an alternative approach, *flg-*Δ3 OFF bacteria from day 9 fecal samples were cultured on TCCFA, then *C. difficile* growth was pooled and tested for swimming motility. No motility was observed, suggesting that the flagellar switch remained in the OFF orientation and that no motile suppressor mutants arose during infection (data not shown). Together these results indicate that preventing flagellar switch inversion impacts *C. difficile* colonization and disease symptom development, with the locked-ON state resulting in greater disease (weight loss) and maintenance of colonization.

### Differences in weight loss is attributable to a higher accumulation of toxins in *flg*-Δ3 ON infected mice

Differences in weight loss between mice infected with the *flg-*Δ3 ON compared to wildtype and *flg-*Δ3 OFF are consistent with the differences in toxin production by these strains. However, the weight recovery observed in mice infected with *flg-*Δ3 ON indicated that this strain may have decreased toxin production through a SigD-independent mechanism. To evaluate this possibility, we determined the toxin titers in fecal samples collected over the 10-day experiments using a Vero cell rounding assay (44). All samples from infected animals caused detectable cell rounding, while no cell rounding occurred when treated with samples from mock-inoculated animals. On day 2, when the differences in weight loss between groups are greatest, the toxin titers in feces collected from *flg*-Δ3 ON-infected mice were significantly higher than from mice infected with wildtype (p = 0.0041); toxin titers were also higher than in feces from *flg*-Δ3 OFF-infected mice, though the differences did not reach statistical significance (Figure 4E). No significant differences in toxin titers between groups of animals were observed for days 4 and 6. On day 8, the *flg*-Δ3 ON samples again had toxin titers higher compared to the wildtype samples (p < 0.05) (Figure 4E). These results suggest that the differences in weight loss observed on day 2 p.i., when bacterial burden was equivalent across groups of infected animals (Figure 4C), is attributable at least in part to a higher accumulation of toxins in mice infected with *flg*-Δ3 ON. However, the higher toxin levels in *flg*-Δ3 ON-infected mice at later stages are likely due to the higher bacterial burden for this strain.

## Discussion

In this study, we used precisely engineered mutations to the right inverted repeat to restrict flagellar switch inversion in *C. difficile* R20291, which allowed us to determine the role of flagellum and toxin phase variation in *C. difficile* physiology *in vitro* and during infection in two rodent models. We characterized mutants with varying abilities to undergo flagellar switch inversion and therefore flagellum and toxin phase variation. *In vitro*, these mutants were either attenuated or fully genotypically and phenotypically locked for swimming motility and toxin biosynthesis but were indistinguishable from wild-type bacteria in growth, sporulation, and germination rates. We analyzed phase-locked mutants in hamster and mouse models of *C. difficile* infection, with distinct outcomes. In hamsters, while the phase-locked mutants led to accumulation of significantly different levels of toxins in the cecum, these differences did not impact acute disease development. In contrast, in the mouse model the mutant with the flagellar switch locked in the ON state caused greater weight loss and persisted longer than wildtype and the locked OFF mutant, which is attributable to higher toxin levels produced by the locked-ON strain in vivo. These findings indicate that the ability to undergo flagellar switch inversion impacts *C. difficile* colonization and disease development.

Previous studies showed that the site specific recombinase RecV is required for inversion of the *flg, cwpV, cmrRST*, and CDR20291_0963 switches, and over-expression of *recV* influences the inversion of the CDR20291_0685 and CDR20291_1514 switches.^24, 42^ Interestingly, the inverted repeats for a given invertible sequence vary in length and in position relative to the determined site of DNA recombination.^24^ Further, sequence conservation among the inverted repeats of RecV-invertible sequences is modest, making the identification of a core RecV-binding sequence difficult. We found that substitution of three residues that are conserved among most of the RIRs (*flg*-3sub) reduced flagellar switch inversion from ON to OFF and OFF to ON but did not eliminate inversion. However, deleting the residue previously determined to be the site of flagellar switch inversion in the RIR and the two flanking residues (*flg*-Δ3), prevented inversion as effectively as deleting the RIR. These results validate the approach of identifying the recombination site by evaluating enrichment of 5’ end clipped reads generated by whole genome sequencing.^24^ Future work will determine whether changes to the site of recombination impair RecV binding and/or the ability to catalyze DNA inversion.

The *C. difficile flg* RIR mutants served to assess the role of flagellar switch inversion on phase variation *in vitro* and *in vivo*. The *flg*-Δ3 and *flg*-ΔRIR mutations resulted in genetically and phenotypically locked strains. The respective ON mutants were motile while the OFF mutants remained non-motile. Consistent with the previously characterized link between flagellum and toxin gene expression via SigD, the ON mutants produced significantly more toxins *in vitro* than the OFF mutants, which produced toxins at a level comparable to the *sigD* control. In contrast to these RIR mutations, the *flg*-3sub mutation appeared to reduce but not eliminate flagellar switch inversion – the inversion assay using *E. coli* showed inversion levels intermediate between the WT and the *flg*-Δ3/*flg*-ΔRIR sequences. In a soft agar swimming motility assay, the *flg*-3sub OFF mutant exhibited motility, possibly because this assay imposes a strong selective pressure for bacteria that can swim to access nutrients;^41^ the *flg*-3sub OFF bacteria that phase varied to *flg* ON would possess an advantage and lead to the observed motile phenotype. Consistent with the swimming motility medium imposing a selective pressure, motile suppressor mutants appeared in some experiments testing the swimming motility of the *flg-*ΔRIR and *flg-*Δ3 OFF mutants, as we observed previously for the *recV flg* OFF mutant.^41^ Unlike in the motility assays, no difference in toxin level was apparent between the *flg*-3sub OFF and the *flg-*ΔRIR and *flg-*Δ3 OFF mutants, likely because the growth conditions for the toxin experiments did not present a selective pressure for the *flg* ON variants.

Prior work by Aubry et al. using the hamster model of acute CDI showed that mutation of genes in the *flgB* operon in *C. difficile* 630Δ*erm* resulted in reduced toxin gene expression, reduced toxin production, and attenuated virulence in hamsters.^32^ Our results showing no discernable difference in CDI development in hamsters between R20291 phase-locked ON and OFF mutants were therefore unexpected. The discrepancy in results may be attributable to strain background. We used 027 ribotype *C. difficile* R20291, which expresses toxin genes and produces toxins at higher levels than the 630 lineage.^36, 47^ There are known differences in the flagellar loci in 630 and R20291 strains,^36^ and we previously showed that *C. difficile* 630 is not capable of flagellum and toxin phase variation.^25^ In addition, mutation of flagellar genes had different effects in R20291 and 630Δ*erm*, a derivative of 630.^32, 33, 48^ We suspect that, despite exhibiting ∼6-fold reduced toxin accumulation compared to the other strains, *flg-*Δ3 OFF nonetheless secreted sufficient toxin to cause acute disease in hamsters given their high sensitivity to *C. difficile* toxins.

Flagella have been shown to play a contributing role in R20291 colonization of the mouse intestinal tract. Non-motile R20291 *fliC* (flagellin), *fliD* (flagellar cap), or *flgE* (hook protein) mutants showed reduced adherence to Caco-2 intestinal epithelial cells *in vitro*, and in a co-infection with wild-type R20291, the *fliC* mutant colonized mice in fewer numbers.^33^ In the current study, the mouse model was more effective at revealing differences between *flg-*Δ3 ON and *flg-*Δ3 OFF. During peak colonization on days 1-3 p.i., *flg-*Δ3 ON resulted in the greatest weight loss, consistent with higher toxin production by this mutant *in vitro*; *flg-*Δ3 OFF did not elicit weight loss in mice, despite being present in equivalent numbers at these time points and producing toxin levels similar to wildtype. During later stages of infection, most mice began recovering weight between days 3 and 4 p.i., when bacterial loads begin to decline, and weights were indiscernible from mock-inoculated animals by day 6. Interestingly, mice infected with *flg-*Δ3 ON recovered their starting weight, even though their numbers were 2-logs higher than wildtype and *flg-*Δ3 OFF. One possible explanation for weight recovery in *flg-*Δ3 ON-infected mice is a SigD-independent (and therefore phase variation independent) down-regulation of toxin production. For example, another toxin gene regulator such as CcpA or CodY could limit toxin synthesis, and SigH, Spo0A, RstA, and SinR could also unlink flagellar and toxin gene expression.^49-53^. However, mean toxin titers remained consistent in feces from *flg-*Δ3 ON-infected mice (compared to decreasing titers from wildtype and *flg-*Δ3 OFF-infected mice). The lack of weight loss in *flg-*Δ3 OFF-infected mice at early time points and in *flg-*Δ3 ON-infected mice at later stages could be due to altered localization of these phase-locked mutants in the intestinal tract, which we speculate results in inefficient delivery of toxins to the epithelium. Both flagellin and the toxins of *C. difficile* have marked immunostimulatory properties and exhibit cooperativity in eliciting an inflammatory response,^18, 54-56^ which may alter the spatial and temporal dynamics of colonization for a strain with constitutively elevated (*flg-*Δ3 ON) or reduced (*flg-*Δ3 OFF) levels of these factors. It is also possible that the observed phenotypes are driven by other mechanisms. SigD, encoded in the *flgB* operon and regulated by the flagellar switch, also regulates genes involved in membrane transport, metabolism, regulation, and cell wall protein synthesis.^39^ These factors may therefore be subject to phase variation and influence *C. difficile* colonization and pathogenesis.

Beyond the population-level analyses of the current study, future work investigating the spatial and temporal dynamics of colonization by individual *flg* ON and *flg* OFF cells, both phase-locked and those arising in a wild-type background, may help clarify the effects of flagellum and toxin phase variation on *C. difficile* colonization and disease development. Does one variant population appear in a particular region of the intestine, either longitudinally or with respect to the epithelium? Does one variant associate with sites of inflammation? Such studies may reveal host and microbiota-derived factors that influence the fitness and virulence of *C. difficile*. Further, because these variants exhibiting different disease potential arise naturally and switch stochastically, flagellum and toxin phase variation may influence not only disease severity but also recurrence. This work may therefore help to identify the *C. difficile* determinants of infection versus asymptomatic carriage which may in turn lead to strategies to distinguish between these potential outcomes, better predict disease severity and recurrence, and mitigate transmission.^57^

## Materials and Methods

### Growth and maintenance of bacterial strains

Strains and plasmids used in this study are listed in Table S1. *C. difficile* strains were grown in an anaerobic chamber (Coy Laboratories) using a gas mix consisting of 85% N_2_, 5% CO_2_, and 10% H_2_. *C. difficile* was routinely cultured in Brain Heart Infusion medium (Becton Dickinson) supplemented with 5% yeast extract (Becton Dickinson) (BHIS) or in Tryptone Yeast (TY) broth as indicated. All *C. difficile* broth cultures were grown at 37°C statically, with 10 μg/mL thiamphenicol (Tm_10_) for plasmid maintenance as needed. *E. coli* DH5α and HB101(pRK24) were cultured under aerobic conditions in LB broth, Miller (Fisher) at 37°C. In *E. coli*, plasmids were maintained with 100 μg/mL ampicillin (Amp_100_), 10 μg/mL chloramphenicol (Cm_10_), and/or 100 μg/mL kanamycin (Kan_100_), as indicated.

### Orientation-specific PCR

*E. coli* strains used in this study are listed in Table S1. Strains used for orientation-specific PCR contain two plasmids: one for expression of *recV* and the other containing the target flagellar switch sequence.^22, 23^ Bacteria were subcultured in BHIS-Cm_10_-Kan_100_ overnight at 37°C and diluted 1:50 into fresh medium. When cultures reached an OD_600_ 0.3-0.4 (early exponential phase), 200 ng/ml anhydrotetracycline (ATc) was added to induce *recV* expression. Cultures were grown until OD_600_ 1.0, and plasmids were purified using the GeneJET Plasmid Miniprep Kit (Thermo Fisher). Purified plasmids were used as template for PCR using primers that discriminate between each flagellar switch sequence orientation. Primers R1614 and R857 were used to amplify the ON orientation of the flagellar switch, which corresponds to the published sequence of R20291 (FN545816.1). Primers R1615 and R857 were used to amplify the OFF orientation of the flagellar switch. All primer sequences are listed in Table S2.

### Generation of mutant strains

To facilitate the generation of mutations in the *flgB* UTR in *C. difficile* R20291, we first deleted the UTR then restored the region with mutant versions of the sequence by allelic exchange with pMSR0, an *E. coli-C. difficile* shuttle vector for toxin-mediated allele exchange mutagenesis.^58^ To delete the *flgB* UTR (*flgB*ΔUTR), upstream and downstream homology regions were amplified from R20291 genomic DNA with R2459 and R2448 or R2449 and R2450, respectively. Gibson assembly was used to introduce these fragments into BamHI-digested pMSR0. Clones were confirmed by PCR and sequencing with plasmid-specific primers R2743 and R2744, which flank the cloning site. The resulting plasmid pRT2546 was introduced into heat-shocked *C. difficile* R20291 via conjugation with *E. coli* HB101(pRK24).^59^ The procedure for allelic exchange was performed as described previously,^58^ except transconjugants were selected and passaged on BHIS-Tm_10_-Kan_100_ agar. Individual colonies were streaked on BHIS-Tm_10_-Kan_100_ agar to ensure purity, then streaked on BHIS-agar with 100 ng/ml ATc (ATc_100_) to induce expression of the toxin-antitoxin genes and eliminate bacteria that still contain pMSR0. Colonies were screened for the desired deletion by PCR with R2451 and R2452. Genomic DNA was isolated from presumptive mutants, amplified with R2451 and R2452, and the resulting PCR product was sequenced with R1512 and R2672 to confirm integrity of the sequence.

Six different inverted repeat mutant constructs were created in *C. difficile flgB*ΔUTR. For all three *flg* ON constructs, RT1702 (*recV flg* ON) genomic DNA was used as the template, and RT1693 (*recV flg* OFF) was used as the template for all three *flg* OFF constructs. Mutants with 3 nucleotide substitutions (*flg-*3sub) were created by changing nucleotides 440-442 of the 498 nt *flgB* UTR from CAA to GTT (Figure 3.2). Mutants with deletions of the right inverted repeat (*flg-*ΔRIR) had nucleotides 424-442 of the *flgB* UTR deleted. A 3 bp deletion (*flg-*Δ3) was made by deleting nucleotides 424-426 of the *flgB* UTR. Overlapping PCR fragments with the desired mutation were amplified and introduced into BamHI/XhoI-digested pMSR0 by Gibson assembly. The following primer pairs were used to amplify the respective fragments (designated as upstream, downstream): *flg-*3sub OFF – R2896 and R2883, R2882 and R2843; *flg-*ΔRIR OFF – R2896 and R2870, R2869 and R2843; *flg-*Δ3 OFF – R2896 and R2885, R2884 and R2843; *flg-*3sub ON – R2896 and R2889, R2888 and R2843; *flg-*ΔRIR ON – R2896 and R2887, R2886 and R2843; *flg-*Δ3 ON – R2896 and R2891, R2890 and R2843. The presence of inserts in pMSR0 was confirmed by PCR with R2743 and R2744, and desired mutations were confirmed by sequencing with R1611, R2313 and R2314. The resulting pMSR0 derivatives were introduced into the *flgB*ΔUTR mutant by conjugation with *E. coli* HB101(pRK24). After selection on BHIS-ATc_100_ plates, colonies were screened with R1512 and R1611 for integration of *flg* constructs. Genomic DNA was isolated from presumptive mutants, amplified with R2451 and R2452, and the resulting PCR product was sequenced with R1512 to confirm integrity of the sequence. The *flg*-Δ3 mutants used in the infection studies and parental R20291 strain were subjected to whole genome sequencing (Microbial Genome Sequencing Center, Pittsburg, PA) to confirm that no unintended sequence polymorphisms arose.

To generate inverted repeat mutants in *E. coli*, the mutant constructs were amplified from the respective pMSR0 plasmids (OFF – *flg-*3sub, *flg-*ΔRIR, *flg-*Δ3; ON – *flg-*3sub, *flg-*ΔRIR, *flg-*Δ3) using R1512 and R1611. These constructs were digested with SphI/EcoRI and ligated into similarly digested pMC123. The presence and integrity of insert was confirmed by PCR with R2313 and R2314 and by sequencing with R2313 and R2462 for *flg* ON and R313 and R2463 for *flg* OFF versions. For inversion assays in *E. coli*, these constructs were co-transformed with pMWO-074::*recV* (ATc-inducible P_*tet*_ promoter) into DH5α.

For overexpression of *recV* in *flg* ON inverted mutant strains, pRT1611 (vector control) or pRT1611::*recV* were introduced by conjugation with *E. coli* HB101(pRK24) and confirmed by PCR.

### Quantitative PCR analysis of the flagellar switch orientation in C. difficile

*C. difficile* strains were grown in BHIS medium to an OD_600_ ∼1.0, and genomic DNA was extracted as previously described.^60^ To analyze flagellar switch orientation from *C. difficile* in mouse feces, fecal samples were suspended in DPBS, treated with lysozyme, and subjected to bead beating to lyse cells including spores. Genomic DNA was purified by phenol/chloroform extraction and washed with ethanol. Quantitative PCR was done using 10 ng (broth culture) or 100 ng (fecal samples) of genomic DNA as template with SYBR Green Real-Time qPCR reagents (Bioline), primers at a final concentration of 1 μM, and an annealing temperature of 55°C. Primers R2175 and R2177 were used to detect the ON orientation and primers R2176 and R2177 were used to detect the OFF orientation. Quantification was done as described previously using the *rpoC* as the control gene.^24^

### Swimming motility assay

For assays using *in vitro* cultures, a single colony from a freshly streaked BHIS plate was inoculated into 0.5X BHIS-0.3% agar to assay flagellum-dependent swimming motility as previously described.^37^ The diameter of motile growth was measured after 48 hours incubation at 37°C. Plates were imaged using the G:BOX Chemi imaging system with the Upper White Light illuminator. To assess swimming motility of *C. difficile* present in mouse feces, diluted fecal samples from each were plated on TCCFA to enrich for *C. difficile*, and growth was pooled and tested for swimming motility in 0.5X BHIS-0.3% agar.

### Detection of TcdA by immunoblot

Immunoblot for TcdA production was performed as previously described.^23, 41, 61^ Cultures were grown overnight (∼16 hours) in TY broth, normalized to an OD_600_ 1.0, and collected by centrifugation at 16,000 x g for 5 minutes. Pellets were suspended in 2x SDS-PAGE sample buffer and boiled for 10 minutes. Samples were electrophoresed on 4%–15% Mini-PROTEAN TGX Precast Protein Gels (Bio-Rad) and transferred to a nitrocellulose membrane (Bio-Rad). Membranes were stained with Ponceau S (Sigma) to assess sample loading and imaged using the G:Box Chemi imaging system. TcdA was detected using mouse α-TcdA antibody (Novus Biologicals) followed by goat anti-mouse IgG secondary antibody conjugated to DyLight 800 4x PEG (Invitrogen). Blots were imaged using the Odyssey imaging system (LI-COR).

### Vero cell rounding assay

We used a previously described protocol to quantify *C. difficile* toxin activity in hamster cecal contents, mouse feces, and bacteria grown overnight in TY broth.^45^ For the assay, 5×10^4^ Vero cells in 90 μl were seeded in each well of a tissue-culture treated, flat bottom 96-well plate (Corning) and allowed to incubate overnight (∼24 hours). The following day, cecal contents collected during necropsy (hamsters) or fecal samples (mice) were thawed at room temperature, weighed, and suspended in 1x DPBS to make an initial 1:10 dilution stock. *In vitro* cultures were grown overnight (∼16 hours) in TY broth, OD_600_ was measured, and cells were pelleted by centrifugation at 16,000 x g for 5 minutes. Supernatants of cecal contents, fecal samples, and broth cultures were sterilized by passing through a 0.45 μm filter. Serial dilutions in 1x DPBS were performed on ice, and 10 μl were applied to each well with Vero cells. DPBS was used as a control. The cells were incubated overnight (∼18 hours) in a tissue culture incubator at 37ºC and an atmosphere with 5% CO_2_, and cell rounding was assessed using a 10x objective on a light microscope. Toxin titer was calculated as the reciprocal of the highest dilution causing ≥80% cell rounding, normalized to OD_600_ of the starting culture (*in vitro*), the amount of cecal contents, or the amount of fecal material (*in vivo*). Samples collected from mock-infected animals were also assayed to show that rounding was specific to *C. difficile*-infected animals.

### Spore purification

Overnight cultures (100 μL) were spread on three to five 70:30 agar plates.^62^ After 72 hours of growth at 37°C, bacterial growth was collected, suspended in 10 mL DPBS and stored aerobically at room temperature overnight. Spores were purified by washing the suspension four times with DPBS before purification using a sucrose gradient as described.^63^ After discarding supernatant that contains cell debris, the spore pellet was washed five more times with DPBS + 1% BSA. Spores were stored in DPBS + 1% BSA at room temperature until use.

### Sporulation assay

Sporulation was assayed as described previously.^64^ Briefly, *C. difficile* strains were grown overnight in BHIS medium supplemented with 0.1% TA and 0.2% fructose to prevent spore accumulation. Cultures were diluted 1:30 in BHIS-0.1% TA-0.2% fructose and upon reaching OD_600_ 0.5, 250 μl of culture was applied to 70:30 agar ^62^. An ethanol resistance sporulation assay was performed at this point to confirm the absence of spores at the initiation of the assay. After 24 hours of growth at 37°C, cells were suspended in BHIS to an OD_600_ 1.0, and an ethanol resistance assay was performed. To eliminate all vegetative cells, 0.5 mL of culture was mixed with 0.5 mL of 57% ethanol to achieve a final concentration of 28.5% ethanol, vortexed and incubated for 15 minutes. To enumerate spores, serial dilutions were made in PBS-0.1% TA and plated on BHIS-0.1% TA agar. To enumerate vegetative cells, serial dilutions of the BHIS cell suspension were plated on BHIS agar. Sporulation efficiency was calculated as the total number of spores divided by the total number of viable cells (spores plus vegetative).

### Germination assay

Spore germination was analyzed at room temperature (27°C) by measuring the change in OD_600_ as previously described.^65^ The germination assay was performed in clear 96-well flat bottom plates (Corning) in a final reaction volume of 100 μl in buffer with 30 mM glycine, 50 mM Tris, 100 mM NaCl, pH 7.5. Spores were suspended in assay buffer, heated at 65°C for 30 minutes, placed on ice for 1 minute and added to wells to a final OD_600_ 0.7. At the start of the assay, 10 mM sodium taurocholate (Sigma Aldrich) (TA) was added to induce germination; no-taurocholate controls were done in parallel. Optical density was measured every 2 minutes for 1 hour using a BioTek Synergy plate reader.

### Ethics statement

Mouse and hamster experiments were performed under the guidance of veterinary staff within the University of North Carolina Chapel Hill Division of Comparative Medicine (DCM). All animal studies were done with prior approval from UNC-CH Institutional Animal Care and Use Committee. Animals that were considered moribund were euthanized by CO_2_ asphyxiation and thoracotomy in accordance with the Panel on Euthanasia of the American Veterinary Medical Association.

### Animal Experiments

Hamster Experiments: Male and female six- to ten-week-old Syrian golden hamsters (Charles River Laboratories) were housed individually and given a standard rodent diet and water *ab libitum*. To induce susceptibility to *C. difficile* infection, one dose of clindamycin (30 mg/kg of body weight) was administered by oral gavage 5 days prior to inoculation. Hamsters were inoculated by oral gavage with approximately 1,000 spores of a single strain of *C. difficile*. Mock inoculated animals were included in each experiment. Fecal samples were collected daily to examine bacterial burden. The animals were monitored at least daily for disease symptoms including weight loss, diarrhea, wet tail, and lethargy. Hamsters that lost 15% or more of their weight or showed severe signs of disease were euthanized by CO_2_ asphyxiation and thoracotomy. Immediately following euthanasia, a necropsy was performed, and cecal contents (not cecal tissue) were collected for enumeration of bacterial CFU, genomic DNA isolation for PCR, and toxin quantification. To enumerate CFU from fecal and cecal contents, samples were weighed, suspended in 1 mL DPBS, heated at 55°C for 20 minutes, and dilutions plated on TCCFA.^66, 67^ To enumerate CFU from the distal colon, the last 3 cm of the colon were excised during necropsy. The samples were weighed, suspended in DPBS, processed with a tissue homogenizer, heated, and plated similarly to cecal and fecal samples. *C. difficile* CFU were enumerated after 48 hours of incubation. Twelve animals (6 male, 6 female) per *C. difficile* strain were tested in two independent experiments. Log rank test for trend was used for statistical analysis of survival data.

#### Mouse experiments

Groups of male and female C57BL/6 mice (Charles River Laboratories) aged 8- to 10-weeks were subjected to a previously described antibiotic regimen to render them susceptible to *C. difficile* infection.^46, 68^ Mice were given a cocktail of kanamycin (400 μg/ml), gentamicin (35 μg/ml), colistin (850 units/ml), vancomycin (45 μg/ml), and metronidazole (215 μg/ml) in their water *ad libitum* seven days prior to inoculation for three days, then returned to regular water for the duration of the experiments. A single intra-peritoneal dose of clindamycin (10 mg/kg body weight) was administered 2 days prior to inoculation. Mice were randomly assigned into groups, with two mice assigned to the mock condition and six mice (3 male, 3 female) to each infection condition. The experiment was independently repeated, and the data were combined for a total of 12 mice (6 male, 6 female) in each infection condition. Mice were inoculated with 10^5^ spores by oral gavage. Mock-inoculated animals were included as controls. Cage changes were performed every 48 h post-inoculation.^68, 69^ Animal weights were recorded, and fecal samples were collected every 24 h for seven days post-inoculation. Fecal samples were homogenized, and dilutions were plated on TCCFA plates, which contain 0.1% of the germinant taurocholate to enumerate spores as colony forming units (CFU) per gram of feces. Additional fecal samples were collected for OS-qPCR analysis to determine flagellar switch orientation and for Vero cell rounding assays to quantify toxins.

## Statistical Analysis and Data Availability

All experiments reflect at least three independent biological replicates, except for animal experiment, which were done twice. Statistical analysis was done using GraphPad Prism 9.1.0. Upon publication, the data that support the findings of this study are available from the corresponding author, R.T., upon reasonable request.

## Acknowledgements

The authors declare no competing interests for this study. This work was supported by NIH award R01-AI107029 and R01-AI143638 to R.T. CLW is supported by an Institutional Research and Academic Career Development Awards (IRACDA) fellowship under K12-GM000678. The funding agency had no role in the design or execution of this study, the analysis of the data, or the decision to publish the results.

## Supporting information

**Table S1. Strains and plasmids used in this study.**

**Table S2. Oligonucleotides used in this study.**

**Figure S1. Mutations in *flg* RIR do not affect growth**. Growth curves of WT, *flg* ON and OFF RIR (3sub, ΔRIR, Δ3) mutants. Overnight cultures grown in TY medium were diluted 1:50 into BHIS broth. Optical density (OD_600_) was measured every 30 minutes for 8 hours. Shown are the means and standard deviation for 6 biological replicates.

**Figure S2. Mutations in *flg* RIR do not affect germination**. Purified spores of indicated strains were germinated in the presence of taurocholate (+) or in buffer without germinant as a control (-), and optical density (OD_600_) was measured. Germination was plotted as the ratio of optical density (OD_600_) at a given time point (t_x_) versus initial OD_600_ (t_0_). A representative germination plot of six independent experiments each consisting of 2 technical replicates is shown.

**Figure S3. Mutations in *flg* RIR do not affect sporulation**. Sporulation efficiency was evaluated by ethanol resistance and calculated as the total number of spores divided by the total number of viable cells (spores plus vegetative). A sporulation-deficient *spo0A* mutant was included as a control. The means and standard deviation of three independent experiments are shown. n.d. – not detectable.

**Figure S4. Bacterial load in feces collected from mice inoculated with WT, *flg*-Δ3 ON, and *flg*-Δ3 OFF**. CFU enumerated in fecal samples collected every 24 hours p.i., with data in Figure 4A separated by day. Day 6 data are shown in Figure 4B. Two independent experiments with n = 6 (3 male, 3 female) were done and the data combined for n = 12. Bars indicate the medians, and dotted lines represent the limit of detection. No CFU were detected in feces of mock-inoculated mice. \*\**p<*0.01, \*\*\**p*<0.001, by Kruskal-Wallis test with Dunn’s post-test comparing all strains.

**Figure S5. The *flg*-Δ3 ON mutant elicits greater weight loss in mice**. Animal weights measured every 24 hours post-inoculation expressed as a percentage of the mouse’s starting weight at day 0, with data from Figure 4C separated by day; day 2 data are shown in Figure 4D. Two independent experiments with n = 6 (3 male, 3 female) were done and the data combined for n = 12. Symbols represent values from individual animals, and bars indicate the means and standard error. \**p*<0.05, \*\**p<*0.01, \*\*\**p*<0.001 by one-way ANOVA with Tukey’s post-test comparing all strains.

