## Supplemental Data for "Flagellum and toxin phase variation impacts intestinal colonization and disease development in a mouse model of Clostridioides *difficile* infection"

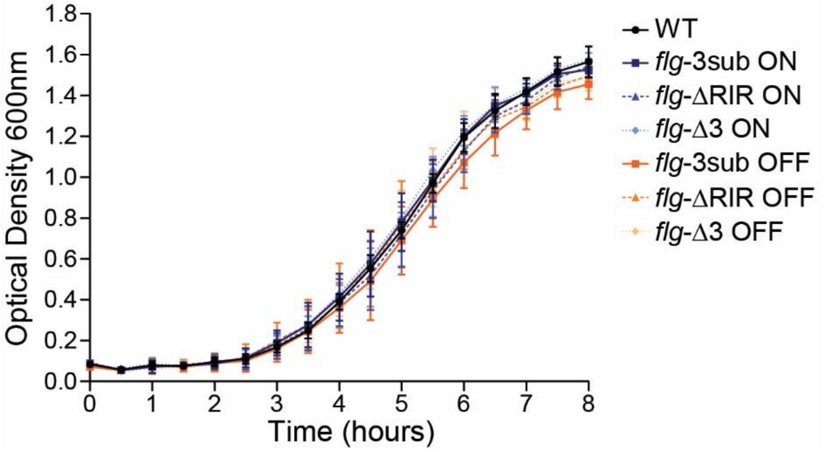


**Figure S1. Mutations in *flg* RIR do not affect growth.** Growth curves of WT, *flg* ON and OFF (3sub, ΔRIR, Δ3) mutants. Overnight cultures grown in TY medium were diluted 1:50 into BHIS broth. Optical density ($OD_{600}$) was measured every 30 minutes for 8 hours. Shown are the means and standard deviations for 6 biological replicates.

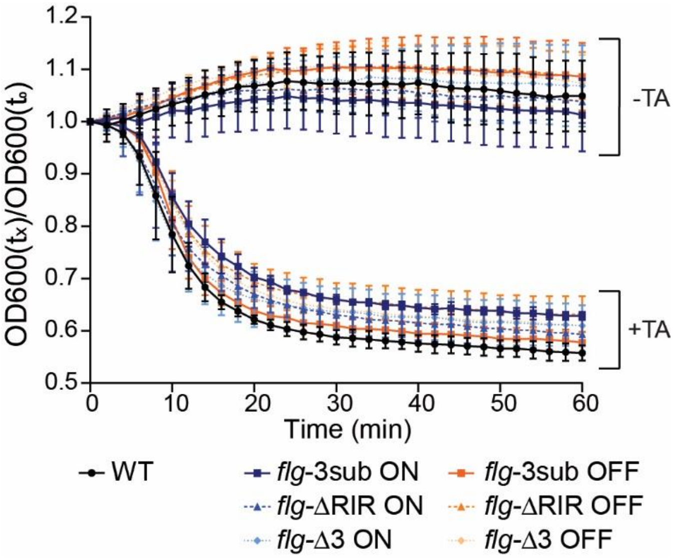


**Figure S2. Mutations in the flagellar switch RIR do not affect germination.** Purified spores of indicated strains were germinated in the presence of taurocholate (+) or in buffer without germinant as a control (-), and optical density ($OD_{600}$) was measured. Germination was plotted as the ratio of optical density ($OD_{600}$) at a given time point ($t_x$) versus initial $OD_{600}$ ($t_0$). A representative germination plot of six independent experiments each consisting of 2 technical replicates is shown. Means and standard deviations are shown.

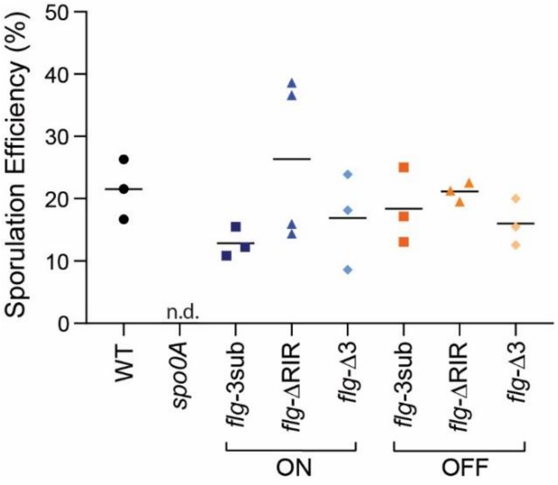


**Figure S3. Mutations in the flagellar switch RIR do not affect sporulation.** Sporulation efficiency was evaluated by ethanol resistance and calculated as the total number of spores divided by the total number of viable cells (spores plus vegetative). A sporulation-deficient *spo0A* mutant was included as a control. The means and standard deviations of three independent experiments are shown. n.d. – none detected. No significant differences by one-way ANOVA other than for the *spo0A* control.

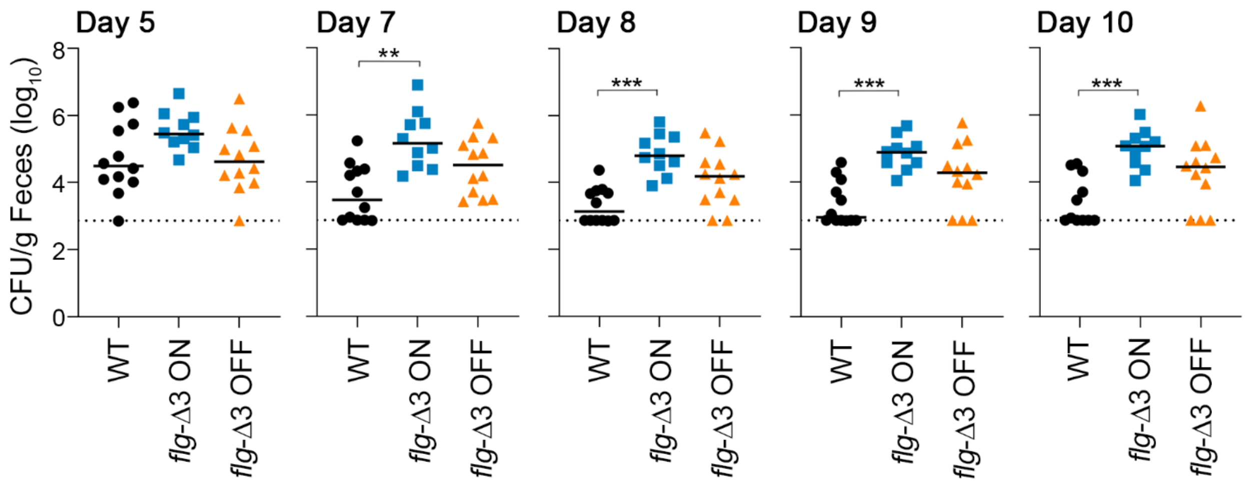


**Figure S4. Bacterial load in feces collected from mice inoculated with WT, *flg*-Δ3 ON, and *flg*-Δ3 OFF.** CFU enumerated in fecal samples collected every 24 hours p.i., with data in Figure 4A separated by day. Day 6 data are shown in Figure 4B. Two independent experiments with n = 6 (3 male, 3 female) were done and the data combined for n = 12. Bars indicate the medians, and dotted lines represent the limit of detection. No CFU were detected in feces of mock-inoculated mice. ***p*<0.01, ****p*<0.001, by Kruskal-Wallis test with Dunn's post-test comparing all strains.

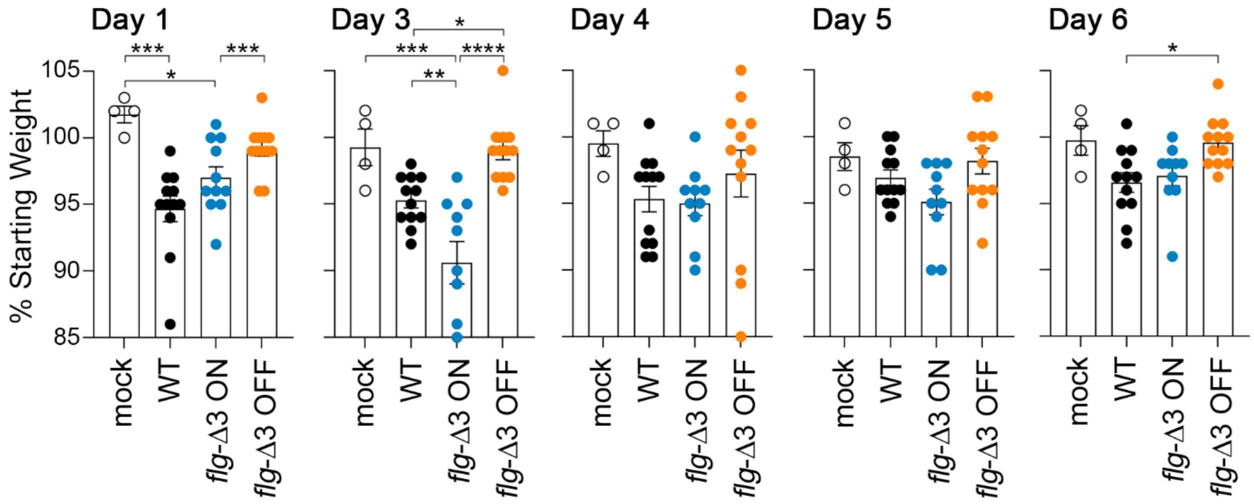


**Figure S5. The *flg*-Δ3 ON mutant elicits increased weight loss in mice.** Animal weights measured every 24 hours post-inoculation expressed as a percentage of the mouse's starting weight at day 0, with data from Figure 4C separated by day; day 2 data are shown in Figure 4D. Two independent experiments with n = 6 (3 male, 3 female) were done and the data combined for n = 12. Symbols represent values from individual animals, and bars indicate the means and standard error. **p*<0.05, ***p*<0.01, ****p*<0.001 by one-way ANOVA with Tukey's post-test comparing all strains.
