## Supplemental Table 1 for "Flagellum and toxin phase variation impacts intestinal colonization and disease development in a mouse model of Clostridioides *difficile* infection"

**Table S1. Strains and plasmids used in this study**

| ***Clostridioides difficile* strains** | | | |
|---|---|---|---|
| **Lab Notation** | **Strain Name** | **Description** | **Reference** |
| RT273 | *C. difficile* R20291 | Ribotype 027 strain | (1) |
| RT1693 | *recV flg* OFF | R20291 *recV::ermB* (*flgB* UTR$^{OFF}$) | (2, 3) |
| RT1702 | *recV flg* ON | R20291 *recV::ermB* (*flgB* UTR$^{ON}$) | (2) |
| RT1566 | *sigD* | R20291 *sigD::ermB* | (2) |
| RT1658 | *spo0A* | R20291 *spo0A::ermB* | (4) |
| RT2555 | *flgB*ΔUTR | R20291 *flgB*ΔUTR | This work |
| RT2603 | *flg*-3sub OFF | R20291 *flg*-3sub OFF | This work |
| RT2604 | *flg*-ΔRIR OFF | R20291 *flg*-ΔRIR OFF | This work |
| RT2609 | *flg*-Δ3 OFF | R20291 *flg*-Δ3 OFF | This work |
| RT2610 | *flg*-3sub ON | R20291 *flg*-3sub ON | This work |
| RT2611 | *flg*-ΔRIR ON | R20291 *flg*-ΔRIR ON | This work |
| RT2612 | *flg*-Δ3 ON | R20291 *flg*-Δ3 ON | This work |
| RT2196 | WT vec | R20291 with pRT1611 (vector control) | (5) |
| RT1562 | WT pRecV | R20291 with pRT1611::*recV* | This work |
| RT2614 | *flg*-3sub ON vec | R20291 *flg*-3sub ON with pRT1611 | This work |
| RT2615 | *flg*-3sub ON pRecV | R20291 *flg*-3sub ON with pRT1611::*recV* | This work |
| RT2616 | *flg*-ΔRIR ON vec | R20291 *flg*-ΔRIR ON with pRT1611 | This work |
| RT2617 | *flg*-ΔRIR ON pRecV | R20291 *flg*-ΔRIR ON with pRT1611::*recV* | This work |
| RT2618 | *flg*-Δ3 ON vec | R20291 *flg*-Δ3 ON with pRT1611 | This work |
| RT2619 | *flg*-Δ3 ON pRecV | R20291 *flg*-Δ3 ON with pRT1611::*recV* | This work |
| ***Escherichia coli* strains** | | | |
| **Lab Notation** | **Strain Name** | **Description** | **Reference** |
| | *Escherichia coli* DH5α | F- φ80*lacZ*ΔM15 Δ(*lacZYA*-*argF*)U169 *recA1 endA1 hsdR*17(rκ -, mκ+) *phoA supE44 thi*-1 *gyrA96 relA1* λ- *tonA* | Invitrogen (6) |
| RT270 | *Escherichia coli* HB101(pRK24) | *E. coli* used in conjugations with *C. difficile*, Amp$^{R}$, Cm$^{R}$ | (7) |
| RT1310 | pFlg OFF pRecV (Ec) | DH5α co-transformed with pRT1324 and pRT1164 | (2) |
| RT2587 | pFlg-3sub OFF pRecV (Ec) | DH5α co-transformed with pRT2582 and pRT1164 | This work |
| RT2588 | pFlg-ΔRIR OFF pRecV (Ec) | DH5α co-transformed with pRT2583 and pRT1164 | This work |
| RT2589 | pFlg-Δ3 OFF pRecV (Ec) | DH5α co-transformed with pRT2584 and pRT1164 | This work |
| RT2602 | pFlg ON pRecV (Ec) | DH5α co-transformed with pRT1323 and pRT1164 | This work |
| RT2590 | pFlg-3sub ON pRecV (Ec) | DH5α co-transformed with pRT2585 and pRT1164 | This work |
| RT2593 | pFlg-ΔRIR ON pRecV (Ec) | DH5α co-transformed with pRT2592 and pRT1164 | This work |
| RT2591 | pFlg-Δ3 ON pRecV (Ec) | DH5α co-transformed with pRT2586 and pRT1164 | This work |

| | | | |
|---|---|---|---|
| **Plasmids** | | | |
| **Lab notation** | **Plasmid Name** | **Description** | **Citation** |
| pRT709 | pRPF185 | pMTL960-derivative, contains ATc-inducible P*tet* promoter | (8) |
| pRT1611 | pRPF185 EV | *gusA* removed from pRPF185 | (2) |
| pRT1529 | pRecV | pRT1611::*recV* | (2) |
| pRT1391 | pMWO-074 | Low copy vector with ATc-inducible P*tet* promoter | (9) |
| pRT1164 | pRecV (Ec) | pMWO-074::*recV* | (2) |
| pRT2460 | pMSR0 | *E. coli-C. difficile* shuttle vector for toxin-mediated allele exchange mutagenesis | (10) |
| pRT2546 | pMSR0 *flgB*ΔUTR | pMSR0 with construct for *flgB* UTR deletion | This work |
| pRT2570 | pMSR0 *flg*-3sub OFF | pMSR0 with construct for *flg*-3sub OFF | This work |
| pRT2571 | pMSR0 *flg*-ΔRIR OFF | pMSR0 with construct for *flg*-ΔRIR OFF | This work |
| pRT2572 | pMSR0 *flg*-Δ3 OFF | pMSR0 with construct for *flg*-Δ3 OFF | This work |
| pRT2573 | pMSR0 *flg*-3sub ON | pMSR0 with construct for *flg*-3sub ON | This work |
| pRT2580 | pMSR0 *flg*-ΔRIR ON | pMSR0 with construct for *flg*-ΔRIR ON | This work |
| pRT2574 | pMSR0 *flg*-Δ3 ON | pMSR0 with construct for *flg*-Δ3 ON | This work |
| pRT264 | pMC123 | pMC-$P_{cpr}$ with nisin-inducible *cpr* promoter | (11) |
| pRT1324 | pFlg OFF | pMC123*::flg* OFF | (2) |
| pRT2582 | pFlg-3sub OFF | pMC123::*flg*-3sub OFF | This work |
| pRT2583 | pFlg-ΔRIR OFF | pMC123::*flg*-ΔRIR OFF | This work |
| pRT2584 | pFlg-Δ3 OFF | pMC123::*flg*-Δ3 OFF | This work |
| pRT1323 | pFlg ON | pMC123*::flg* ON | (2) |
| pRT2585 | pFlg-3sub ON | pMC123::*flg*-3sub ON | This work |
| pRT2592 | pFlg-ΔRIR ON | pMC123::*flg*-ΔRIR ON | This work |
| pRT2586 | pFlg-Δ3 ON | pMC123::*flg*-Δ3 ON | This work |
