## Supplemental Table 2 for "Flagellum and toxin phase variation impacts intestinal colonization and disease development in a mouse model of Clostridioides *difficile* infection"

**Table S2. Oligonucleotides used in this study**

| Lab Notation | Primer name | Sequence (5' to 3')[a] | Reference |
|---|---|---|---|
| R850 | rpoCqF | CTAGCTGCTCCTATGTCTCACATC | [97] |
| R851 | rpoCqR | CCAGTCTCTCCTGGATCAACTA | [97] |
| R2175 | flg_switchON_qF | GTTTTCTTACCAAAGTGATACATTATTATATTAATG | [133] |
| R2176 | flg_switchOFF_qF | CATTAATATAATAATGTATCACTTTGGTAAGAAAAC | [133] |
| R2177 | flg_switch_qR | GCTATTGTCTGACTTCTTAAATTAGTTGCAT | [133] |
| R2313 | flg_switch_F | ATCACATTATGTAGTAAAAAACACC | This work |
| R2314 | flg_switch_R | GTATTCACACTCACTCCTCC | This work |
| R2462 | flgON_RIR_F | ATATTAGTTTTCTTACCAAAGTGATAC | This work |
| R2463 | flgOFF_RIR_F | TAATAATGTATCACTTTGGTAAGAAAAC | This work |
| R857 | flgBqR | AGGCATAGCATCATTTAGTGTTTCTTC | [97] |
| R1614 | CDR20291_0248InvF | AGGCAACTTTATAAAGAAATATTTAAATTTATATTAAAATATTTTTATATTTTTATTAGG | [133] |
| R1615 | CDR20291_0248InvR | CCTAATAAAAATATAAAAATATTTTAATATAAATTTAAATATTTCTTTATAAAGTTGCCT | [133] |
| R2459 | flgB_UTR_F1 | CAGGAAACAGCTATGACCGCGGCCGCGATTGTGCTCAATCTCATGG | This work |
| R2448 | flgB_UTR_R1 | CACACTCACTCCTCCTCACTATTTAGTTTTAACTTAAGTATACAATAAATAAC | This work |
| R2449 | flgB_UTR_F2 | GTTATTTATTGTATACTTAAGTTAAAACTAAATAGTGAGGAGGAGTGAGTGTG | This work |
| R2450 | flgB_UTR_R2 | GATCGCGCATGTCTGCAGGCCTCGAGCTTATCAACTTCTGTTCTAGGTAC | This work |
| R2672 | flgBΔUTR_screen_F | GGAGATGCAGGAGCTATAG | This work |
| R2741 | flgBΔUTR_F1 | CATTGATTTCTTTCAGTTTCGGATCCTTGTGCTCAATCTATGGG | This work |
| R2742 | flgBΔUTR_R2 | GACGTCGACTCTAGAGGATCCCATAGATAGCTGTGCTTCTTGACC | This work |
| R2743 | pMSR0_screen_F | GTGTTATCAATTGCACTACTCATGG | This work |
| R2744 | pMSR0_screen_R | GTTGAACCATTAGCTAAGGATTCAG | This work |
| R2896 | flg_ΔUTR_insertion_F1 | GTGTCCATTGATTTCTTTCAGTTTCGGATCCGATTGTGCTCAATCTCATGGAG | This work |
| R2843 | flg_ΔUTR_insertion_R2 | CTTGCATGTCTGCAGGCCTCGAGATAGCTGTGCTTCTTGAC | This work |
| R2882 | flgOFF_3nt subs_F2 | GTTGCCTTTTTTTGTAATATAGGTTCTTCATTTTTTATTAATAAGC | This work |
| R2883 | flgOFF_3nt subs_R1 | GCTTATTAATAAAAAATGAAGAACCTATATTACAAAAAAAGGCAAC | This work |
| R2869 | flgOFF_ΔRIR_F2 | CTTTATAAAGTTGCCCTTCATTTTTTATTAATAAGC | This work |
| R2870 | flgOFF_ΔRIR_R1 | GCTTATTAATAAAAAATGAAGGGCAACTTTATAAAG | This work |
| R2884 | flgOFF_Δ3nt_F2 | TATTTCTTTATAAAGTTGCCTTTTGTAATATAGCAACTTCATTTTTTATTAATAAG | This work |
| R2885 | flgOFF_Δ3nt_R1 | AAAAAATGAAGTTGCTATATTACAAAAGGCAACTTTATAAAG | This work |
| R2888 | flgON_3nt subs_F2 | GAATAAAGAAGTCATTTTTTGTAATATAGGTTCTTCATTTTTTATTAATAAGC | This work |
| R2889 | flgON_3nt subs_R1 | GCTTATTAATAAAAAATGAAGAACCTATATTACAAAAAATGACTTCTTTATTC | This work |

| | | | |
|---|---|---|---|
| R2886 | flgON_ΔRIR_F2 | GTAATTAATTTGGATGAATAAAGAAGTCCTTC ATTTTTTATTAATAAGC | This work |
| R2887 | flgON_ΔRIR_R1 | GCTTATTAATAAAAAATGAAGGACTTCTTTAT TCATCCAAATTAATTAC | This work |
| R2890 | flgON_Δ3nt_F2 | GTAATTAATTTGGATGAATAAAGAAGTCTTTT GTAATATAGCAACTTC | This work |
| R2891 | flgON_Δ3nt_R1 | TAAAAAATGAAGTTGCTATATTACAAAAGACT TCTTTATTCATCCAAATTAATTAC | This work |
| R1512 | CDR202_PflgBnew | GTTCAAGCATGCGATATATTGTACAAATAAAA TTGAAATATATGG | [133] |
| R1611 | CDR20291_0248CDSR | CAAGAATTCTTACCTCCCACTTATTATTGA | [133] |

[a] Restriction sites used for cloning are underlined
